# The Unlabeled Organ: Adipose tissue and normativity in anatomy and physiology texts

**DOI:** 10.64898/2026.07.14.736897

**Authors:** Isabel D.K. Hermsmeyer, Ormond A. MacDougald, Mary E. Orczykowski

## Abstract

Adipose tissue is a dynamic and essential organ system with roles in metabolism, endocrine signaling, immune function, thermoregulation, and structural support. Despite this, it has historically been framed as inert or pathological, raising questions about how it is represented in biomedical education. We evaluated the visual and textual representation of adipose tissue across widely used anatomy, physiology, and combined anatomy–physiology textbooks. Images were coded for presence, labeling, and degree of representation, and text mentions of “adipose” and “fat” were quantified across functional contexts. In anatomy textbooks, adipose tissue was frequently present (6.6–21.7% of images) but rarely labeled (<5% of all images; <25% of adipose-containing images). Representation was often minimal or incidental, and text descriptions were predominantly non-functional, with a preference for the term “fat.” In physiology textbooks, adipose tissue was more often described in functional terms and more frequently referred to as “adipose,” particularly in metabolic contexts, but appeared in relatively few images (<5%) and was typically minimally depicted. Combined anatomy–physiology texts showed intermediate patterns but retained low labeling rates and limited integration of structure and function. Across all textbook types, terminology varied by context, with “fat” more common in non-functional and obesity-related descriptions and “adipose” associated with metabolic functions, suggesting context-dependent bias in terminology. Overall, adipose tissue is present but inconsistently framed across visual and textual domains. This fragmentation may limit recognition of adipose tissue as an integrated organ system and may reinforce reductive or stigmatizing interpretations. Aligning educational representations with current scientific understanding is therefore essential to support a more accurate and integrated view of adipose tissue as a dynamic organ system.

## Introduction

Adipose tissue is now widely recognized as a dynamic and essential organ system, with roles spanning energy homeostasis, endocrine signaling, immune function, thermoregulation, and physical support (Emont & Rosen; 2023; Grant & Dixit; 2015; Kahn et al., 2019; Loft et al., 2025; Luong et al., 2019). Historically, however, adipose tissue has often been conceptualized as a relatively inert or excess energy store (Corvera, 2021a; Cypess, 2022). In anatomical contexts, it is frequently treated as background material, and removed during dissection to facilitate visualization of other structures, while in clinical and educational settings it is often primarily discussed in relation to metabolic disease (Goss et al., 2020; Harvey et al., 2020; Lafontan, 2012; Richard et al., 2020). Together, these framings position adipose tissue as incidental or pathological rather than as a structurally organized and physiologically essential component of the human body.

The importance of adipose tissue is perhaps most apparent when its quantity or function is disrupted. Excess adiposity is associated with insulin resistance, type 2 diabetes, cardiovascular disease, cancer, and numerous other chronic conditions, whereas severe adipose deficiency or pathological loss results in profound metabolic and endocrine dysfunction, immune perturbation, ectopic lipid deposition, and structural complications that arise when supporting depots are depleted (Daley et al., 2025; Maung et al., 2025; Miricescu et al., 2021; Sahni et al., 2017). In severe cases, both excess and deficiency can be life-threatening (Blüher, 2019; Maung et al., 2026). These opposing clinical outcomes illustrate that adipose tissue is not merely an expendable energy reserve but is critical for maintaining physiological homeostasis.

This functional importance is reflected in its highly organized anatomy. Rather than existing as a diffuse or passive accumulation of lipid, adipose tissue is organized into anatomically distinct depots distributed throughout the body (Bagchi & MacDougald, 2019; Zwick et al., 2018). These depots are commonly classified into either subcutaneous or visceral categories, although numerous anatomically localized depots fall outside this framework. Subcutaneous adipose tissue lies beneath the skin, whereas visceral adipose tissue is distributed around internal organs within the thoracic, abdominal, and pelvic cavities (Ibrahim, 2010; Zwick et al., 2018). While both contribute to whole-body metabolic and endocrine homeostasis, visceral depots are generally more metabolically and immunologically active (Fox et al., 2007; Macdougall et al., 2018). Importantly, both subcutaneous and visceral adipose tissues comprise multiple anatomically distinct depots with unique developmental origins, vascular supply, and physiological characteristics. Numerous additional adipose depots are distributed throughout the body, including but not limited to bone marrow and intra-articular adipose tissues, all characterized by unique anatomical locations and physiological roles (Loft et al., 2025; Zwick et al., 2018).

Adipose tissue is equally heterogeneous at the cellular level. White adipose tissue, composed predominantly of large unilocular lipid-filled adipocytes, serves as the body’s principal site of energy storage while also functioning as a major endocrine and immune organ (Trayhurn & Beattie, 2001). In contrast, brown adipose tissue contains multilocular adipocytes enriched with mitochondria expressing uncoupling protein-1 (UCP1), enabling the dissipation of chemical energy as heat through non-shivering thermogenesis. Although most abundant during infancy, brown adipose tissue persists in smaller quantities in many adults. Beige adipocytes represent an inducible thermogenic cell type that emerges within white adipose depots in response to environmental and physiological stimuli, acquiring characteristics similar to brown adipocytes (Cypess, 2022; Devlin; 2021). Adipocytes themselves represent fewer than half of the total cellular content of adipose tissue despite comprising most of its volume. The remaining cellular population includes adipose progenitor cells, vascular cells, and diverse innate and adaptive immune cells that are essential for adipose tissue development, maintenance, and function (Corvera, 2021b). Cellular specialization also exists within individual depots. For example, bone marrow adipose tissue comprises constitutive and regulated subtypes that differ in anatomical distribution, metabolic responsiveness, and interactions with hematopoiesis and skeletal homeostasis (Scheller et al., 2015; Li et al., 2022). Finally, adipocytes are adapted to the distinct thermal environments of different depots. Peripheral depots, including subcutaneous and distal bone marrow adipose tissues, generally exist at lower temperatures than visceral depots within the body core, promoting depot-specific metabolic adaptations (Mori et al., 2021).

Adipose tissue has physiological roles that extend well beyond energy storage. In addition to serving as the body’s major energy reservoir, it helps regulate systemic lipid and glucose homeostasis through the uptake, storage, and mobilization of nutrients (Frühbeck et al., 2001; Tseng, 2023). Endocrinologically, adipocytes secrete numerous bioactive molecules, including leptin, adiponectin, and other adipokines, that influence appetite, reproduction, bone metabolism, insulin sensitivity, and energy expenditure (Ahima & Flier, 2000; Kershaw & Flier, 2004). Adipose tissue also functions as an immunological organ, housing diverse innate and adaptive immune cell populations that contribute to tissue homeostasis and inflammatory responses (Grant & Dixit, 2015; Macdougall et al., 2018). In brown and beige adipose tissues, specialized thermogenic adipocytes dissipate chemical energy as heat to support body temperature regulation (Cypess, 2022). Across the body, adipose tissue also provides essential structural functions, including mechanical cushioning, shock absorption, and physical support for joints, neurovascular structures, and internal organs (Dragoo et al., 2021; Labusca & Zugun-Eloae, 2018; Wronska & Kmiec, 2012). Collectively, this multilayered anatomical, cellular, and functional heterogeneity establishes adipose tissue as a highly integrated organ system rather than a homogeneous tissue.

Despite this, the extent to which adipose tissue is represented in foundational educational materials remains unclear. Prior work has suggested that adipose tissue is minimally depicted, if at all, in anatomical atlases (Cypess, 2022), but this claim has not been systematically evaluated. Observations from anatomy instruction further suggest that adipose tissue is frequently presented as an obstacle to dissection rather than as a structure of interest in its own right (Goss et al., 2020). Much of the work redefining adipose tissue as a metabolically active and functionally diverse organ system has emerged from physiological research (Ahima & Flier, 2000; Lafontan, 2012; Zwick et al., 2018), raising the question of whether these advances are reflected in physiology educational materials and how this compares to its apparent absence or de-emphasis in anatomical contexts. Accordingly, these disciplinary differences may contribute to fragmented representations of adipose tissue across educational materials, with implications for how it is conceptualized by learners.

Understanding how adipose tissue is represented across educational resources is important, as these materials shape how students conceptualize the body and its component systems. Incomplete or selective representation of adipose tissue may limit recognition of its full biological and clinical significance. While adipose tissue is often emphasized at the extremes, from obesity to lipodystrophy and cachexia, its importance extends across the full spectrum of physiological states (Daley et al., 2025; Dragoo et al., 2021; Maung et al., 2026). Variation in adipose distribution and function, even within clinically normal ranges, influences metabolic health, endocrine signaling, and disease risk, underscoring its relevance in both health and pathology. A thorough understanding of its physiological functions must be paired with anatomical literacy; recognizing where adipose tissue is distributed, how it is organized into distinct depots, and how it interfaces with other organ systems. Without this spatial framework, adipose tissue may be misinterpreted or overlooked, with consequences for imaging interpretation, surgical approaches, and the recognition of normal anatomical variation.

When educational materials fail to present adipose tissue as a structured and functionally essential organ system, students may develop incomplete frameworks for understanding both health and disease (Goss et al., 2020; Nutter et al., 2023; Puhl, 2023). In such contexts, interpretation of adipose tissue may rely more heavily on prevailing societal narratives, which often frame “fat” primarily as pathological or excessive, rather than on its numerous scientifically established physiological roles (Harleen et al., 2026; Heidebrecht et al., 2024). This framing is not neutral: weight stigma is well documented in medical education and clinical practice and can influence both provider attitudes and patient care (Puhl, 2023). Within this context, larger-bodied individuals with higher adiposity are often assumed to be less healthy, even in the absence of clinical evidence (Goss et al., 2020), a bias that is further reflected in patterns of representation within educational materials, where larger-bodied individuals are rarely depicted in anatomical imagery (Beresheim et al., 2024). As a result, while all individuals can be affected by incomplete representation, larger-bodied individuals are more likely to have their physiology interpreted primarily through a pathological lens, with potential consequences for clinical assessment, decision-making, and care.

In this study, we systematically evaluate the representation of adipose tissue in widely used anatomy, physiology, and combined anatomy and physiology textbooks. We quantify both visual and textual depictions of adipose tissue, characterize how it is described across contexts, and compare patterns of representation between disciplines. By doing so, we aim to provide an empirical foundation for assessing how adipose tissue is positioned within biomedical education and to identify opportunities for more accurate and inclusive representation.

## Methods

### Textbook selection

Textbooks were selected to represent commonly used resources in anatomy, physiology, and combined anatomy–physiology education at both undergraduate and graduate levels (See Table 1). Candidate textbooks were identified through consultation with anatomy and physiology faculty, review of course syllabi from undergraduate and medical/graduate-level courses, and searches of widely adopted anatomy and physiology textbooks used in higher education. Only the most up-to-date edition of each text was included in the coded analyses. While not all courses rely on a single textbook, and some courses may not use textbooks at all, these texts represent widely used instructional resources and reflect current common approaches to teaching anatomy and physiology. The final sample was intended to capture a broad cross-section of instructional materials commonly encountered in anatomy and physiology education. All textbooks included in the final sample were subjected to both visual (image-based) and textual coding, allowing direct comparison of adipose tissue representation across visual and textual domains throughout.

**Table 1.** Adipose tissue representation in anatomy textbook images. Texts are listed in descending order from highest proportion of visual adipose tissue representation to the lowest. Percentages are calculated relative to the total number of images within each textbook.

| Text | Edition | Year | Type | Total images | Adipose images, n (%) | Images with adipose labels, n (%) | Depot-specific labels, n (%) |
| --- | --- | --- | --- | --- | --- | --- | --- |
| Netter Atlas of Human Anatomy | 8th | 2023 | Atlas | 1224 | 265 (21.7%) | 50 (4.1%) | 25 (2%) |
| Rohen's Photographic Anatomy | 9th | 2021 | Atlas | 1188 | 245 (20.6%) | 11 (0.9%) | 2 (0.2%) |
| Grant's Atlas of Anatomy | 16th | 2017 | Atlas | 1869 | 343 (18.4%) | 74 (4%) | 36 (1.9%) |
| Gosling's Human Anatomy: Color Atlas and Textbook | 6th | 2017 | Atlas | 641 | 102 (15.9%) | 20 (3.1%) | 13 (2%) |
| Gilroy Atlas of Anatomy | 5th | 2025 | Atlas | 1939 | 242 (12.5%) | 40 (2.1%) | 31 (1.6%) |
| Sobotta Atlas of Human Anatomy | 16th | 2019 | Atlas | 1077 | 132 (12.3%) | 12 (1.1%) | 3 (0.3%) |
| Moore's Clinically Oriented Anatomy | 10th | 2026 | Clinical Textbook | 2416 | 253 (10.5%) | 36 (1.5%) | 14 (0.6%) |
| The Big Picture: Gross Anatomy | 2nd | 2019 | Textbook | 411 | 43 (10.5%) | 13 (3.2%) | 9 (2.2%) |
| Tortora's Principles of Anatomy | 15th | 2020 | Textbook | 1462 | 125 (8.6%) | 17 (1.2%) | 10 (0.7%) |
| Gray's Anatomy: The Anatomical Basis of Clinical Practice | 43rd | 2026 | Textbook | 4089 | 349 (8.5%) | 51 (1.2%) | 27 (0.7%) |
| Gray's Anatomy for Students | 5th | 2023 | Textbook | 2320 | 172 (7.4%) | 29 (1.3%) | 18 (0.8%) |
| Abrahams' and McMinn's Clinical Atlas of Human Anatomy | 8th | 2020 | Clinical Atlas | 1658 | 116 (7%) | 27 (1.6%) | 19 (1.2%) |
| Marieb's Human Anatomy | 9th | 2020 | Textbook | 1210 | 80 (6.6%) | 20 (1.7%) | 3 (0.3%) |

### Image Coding

All images within each textbook were counted to determine the total number of figures per text. The unit of analysis for visual content was the individual figure panel; multi-panel figures were coded separately for each panel when panels contained distinct images. Repeated images appearing multiple times within a textbook were counted each time they appeared, as they represent repeated exposure to the image in the instructional material.

Images were reviewed individually to identify those in which adipose tissue was visually present. In photographic images, adipose tissue was identified by its typical lobulated appearance and coloration consistent with adipose tissue in anatomical dissections or surgical images, recognizing that coloration may vary depending on tissue condition (e.g., living tissue, surgical exposure, or embalmed donor tissue). In illustrated figures, adipose tissue was identified by yellow, globular structures corresponding to adipose lobules. Artistic depictions in which yellow globular tissue was illustrated in anatomical locations where adipose tissue does not normally occur were not included in the adipose image count. For example, several illustrations depicted yellow lobular tissue within the shaft of the penis, likely reflecting an accidental conflation of fascial tissues with adipose; these images were excluded from adipose classification.

Only images in which adipose tissue could be visually identified were subjected to further coding. For each image containing adipose tissue, several variables were recorded. These included whether adipose tissue was explicitly labeled in the image and/or in the figure caption, and whether the label specified a particular anatomical depot. Labels were considered adipose-specific if the image explicitly identified adipose tissue or fat. Depot-specific labels were defined as labels identifying a named anatomical adipose depot (e.g., perirenal adipose tissue, epidural adipose tissue, infrapatellar fat pad). References describing adipose tissue simply as occurring within an anatomical space (e.g., adipose tissue in the ischioanal fossa) were not considered depot-specific, as these descriptions indicate adipose tissue incidentally present within a location rather than identifying a distinct anatomical adipose depot.

To assess how adipose tissue was visually depicted, each image containing adipose tissue was assigned a score on a four-point representation scale reflecting the degree to which adipose tissue was visually retained in locations where it would normally occur anatomically. A score of 1 indicated that adipose tissue was largely removed or minimized, such as in simplified “cutaneous cutout” images in which the tissue’s presence is depicted only as a thin band within the hypodermis. A score of 2 indicated that adipose tissue was present but reduced relative to its expected anatomical presence. A score of 3 indicated moderate representation of adipose tissue in anatomically typical locations. A score of 4 indicated that adipose tissue was depicted in a manner consistent with typical anatomical appearance, with adipose retained wherever it would normally be present.

We did not attempt to determine when adipose tissue *should* have been depicted but was absent. Because many images are designed to emphasize specific structures (e.g., skeletal or muscular systems), the expected presence of adipose tissue is inherently context-dependent and difficult to define objectively. As such, analyses were restricted to images in which adipose tissue was visibly present, and representation was quantified based on extent rather than absence.

### Text coding

Textual references to adipose tissue were also analyzed. All textbooks were accessed electronically (e.g., PDF editions or online platforms such as ClinicalKey and VitalSource), allowing full-text keyword searches. All occurrences of the term adipose and its variants (e.g., adipocyte, adiposity) were recorded. Occurrences of the term fat and related variants (e.g., fatty) were recorded only when the term was used linguistically to refer to adipose tissue. Uses of fat referring to other biochemical or nutritional contexts (e.g., fat-soluble vitamins, fat storage in the liver) were not included. Identified references were categorized based on the functional context in which adipose tissue was discussed. Each mention was classified into one of several categories: indirect or non-functional description, obesity or excess, metabolism, thermogenic function, endocrine signaling, structural roles, and immune-related processes.

We did not attempt to quantify adipose tissue representation relative to other tissues, as no clear or consistent baseline for comparison exists across systems-based content. Instead, we quantified absolute mention counts (with page-level tracking) and focused on the contextual framing and terminology used (e.g., “fat” versus “adipose”).

All coding was conducted by a single investigator (*.*.) using a predefined coding protocol. To assess intra-coder reliability, a randomly selected subset of images and text references (10% of the dataset; n = 592) was re-coded after an interval of several weeks. Percent agreement between coding rounds was 94%, indicating high intra-coder reliability.

### Qualitative methods

In addition to the quantitative coding described above, qualitative observations were recorded during the review of both images and text. These included patterns in how adipose tissue was visually depicted, the contexts in which adipose tissue appeared in figures, and the language used to describe adipose tissue in accompanying text. Illustrative examples and representative quotations were collected to characterize recurring narrative and visual trends across textbooks. These qualitative observations were used to contextualize the quantitative findings and to highlight common pedagogical patterns in the representation of adipose tissue.

### Statistical analysis

For anatomy image analyses, adipose representation and labeling were quantified as counts and proportions within each textbook. Differences in labeling rates were evaluated using binomial generalized linear models, with the number of labeled images modeled relative to total adipose images and textbook type included as a predictor. Representation score distributions were summarized as proportions across categories, and differences in distributions were assessed using chi-square tests, followed by pairwise comparisons of proportions with Benjamini–Hochberg adjustment. Confidence intervals for proportions were estimated using binomial tests.

For text-based adipose representation, we quantified the number of mentions of the terms fat and adipose within predefined contextual categories (e.g., obesity/excess, metabolism) across physiology textbooks. Differences in contextual usage were evaluated using binomial mixed-effects models, with context-specific mentions modeled relative to total term mentions, term (fat vs. adipose) included as a fixed effect, and textbook as a random intercept. Effects were expressed as odds ratios (ORs), and p-values were adjusted for multiple comparisons using the Benjamini–Hochberg procedure.

## Results

### Anatomy Textbooks

The proportion of images containing adipose ranged from 6.6% to 21.7% of total images, with atlases generally exhibiting higher proportions than general and clinical textbooks (Table 1). Absolute counts of adipose-containing images were also substantial in several widely used references (Figure 1), including *Gray’s Anatomy: The Anatomical Basis of Clinical Practice* (n = 349), *Grant’s Atlas of Anatomy* (n = 343), and the *Netter Atlas of Human Anatomy* (n = 265). However, the proportion of adipose images that were explicitly labeled varied across textbooks but was consistently low overall (see Table 1, Figure 1C). Across all texts, labeling rates generally fell below 25% of adipose-containing images and below 5% of all images, with several textbooks exhibiting substantially lower rates. Depot-specific labeling was consistently rarer than general adipose labeling, typically comprising a small subset of labeled images. Despite variation among individual textbooks, labeling rates did not differ significantly by textbook type (binomial model, all p > 0.5). Estimated effects were small (clinical vs. atlas OR = 0.91; textbook vs. atlas OR = 1.04), indicating that under-labeling of adipose tissue is consistent across atlases, clinical texts, and general textbooks.

**Figure 1:**
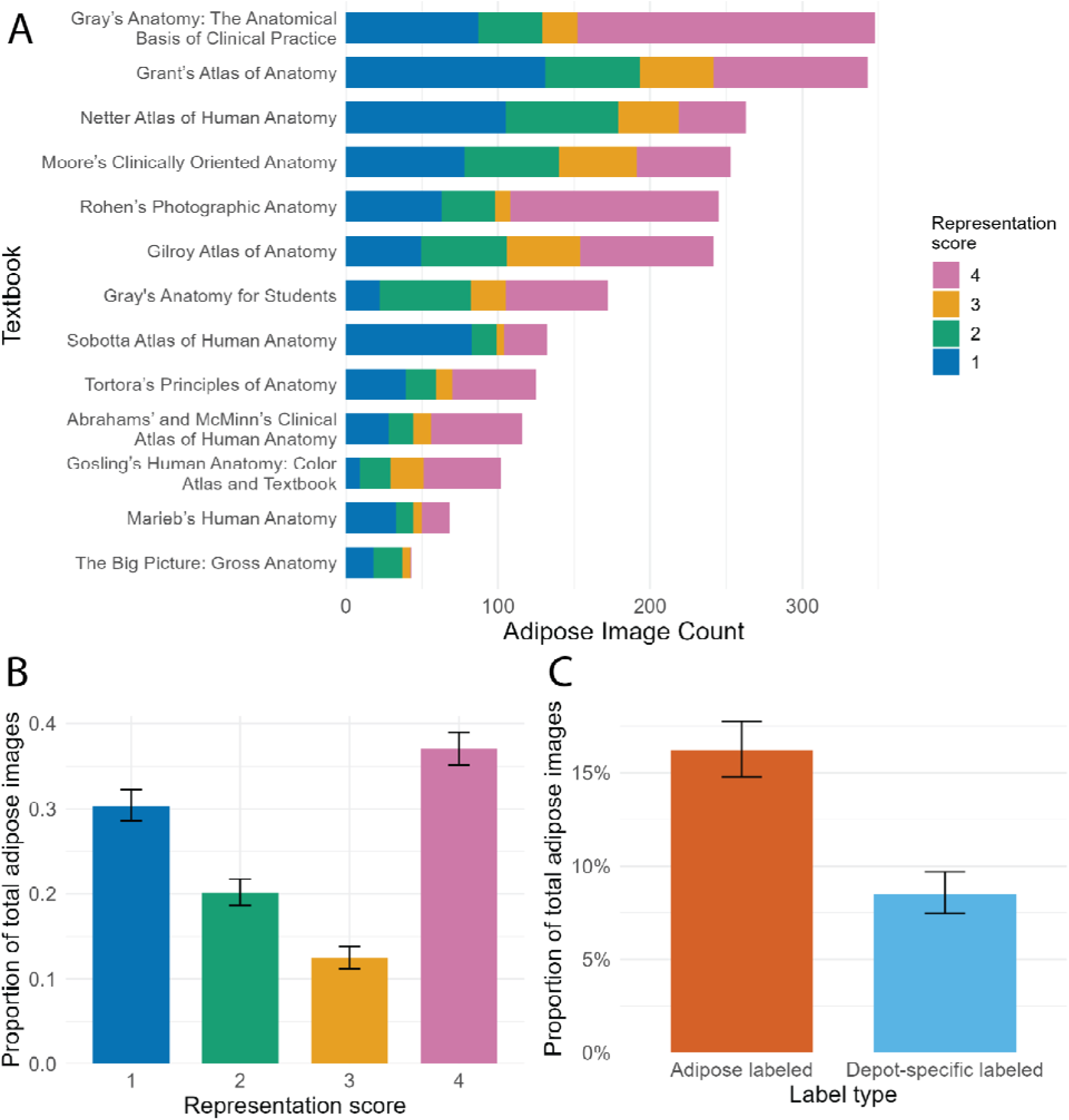
Adipose tissue representation and labeling in anatomy textbook images. (A) Counts of adipose-containing images by textbook, stratified by representation score (1–4), with higher scores indicating more complete depiction. (B) Proportion of adipose images in each representation category (mean ± 95% CI). (C) Proportion of images containing adipose with adipose labeling, including any label and depot-specific labels (mean ± 95% CI).

Adipose images were unevenly distributed across representation score categories (Figure 1B). Representation scores differed significantly from a uniform distribution (χ² = 348.24, df = 3, p < 0.001). The highest representation category (score 4) comprised the largest proportion of images (37.0%, 95% CI: 35.1–39.0), followed by score 1 (30.4%, 95% CI: 28.6–32.2). Lower proportions were observed for intermediate categories (score 2: 20.1%, 95% CI: 18.6–21.8; score 3: 12.4%, 95% CI: 11.2–13.8). Pairwise comparisons indicated that all representation categories differed significantly from one another after correction for multiple comparisons (Benjamini–Hochberg-adjusted p < 0.001 for all comparisons), indicating a structured and non-uniform distribution of representation across score levels.

In-text mentions of adipose tissue and fat were also assessed across anatomy textbooks (Table S1, Figure S1). Across texts, references to “fat” were generally more frequent than references to “adipose,” though total mention counts varied widely by textbook. Most mentions occurred in indirect or non-functional contexts, typically describing where adipose (or fat) is anatomically located on the body. A smaller representation of in text mentions were for structural roles, with relatively fewer references to metabolic, endocrine, or immune roles, as may be typically expected within anatomy (without physiology) texts. Overall, across anatomy texts, in-text mentions predominantly use the term “fat” rather than “adipose” and are largely focused on describing where fat is located on the body.

### Physiology Textbooks

In-text mentions of “adipose” and “fat” varied across physiology textbooks, with total mention counts ranging from 37 to 151 per text (Table 2). In most textbooks, references to “adipose” were as frequent or more frequent than references to “fat,” particularly in widely used physiology texts such as Vander’s Human Physiology and Silverthorn Human Physiology. However, some texts, including Guyton and Hall Textbook of Medical Physiology and Barrett (Ganong) Review of Medical Physiology, exhibited more balanced or higher use of the term “fat.” Overall, both terms were used extensively across texts, with variation in relative frequency by textbook.

**Table 2.**
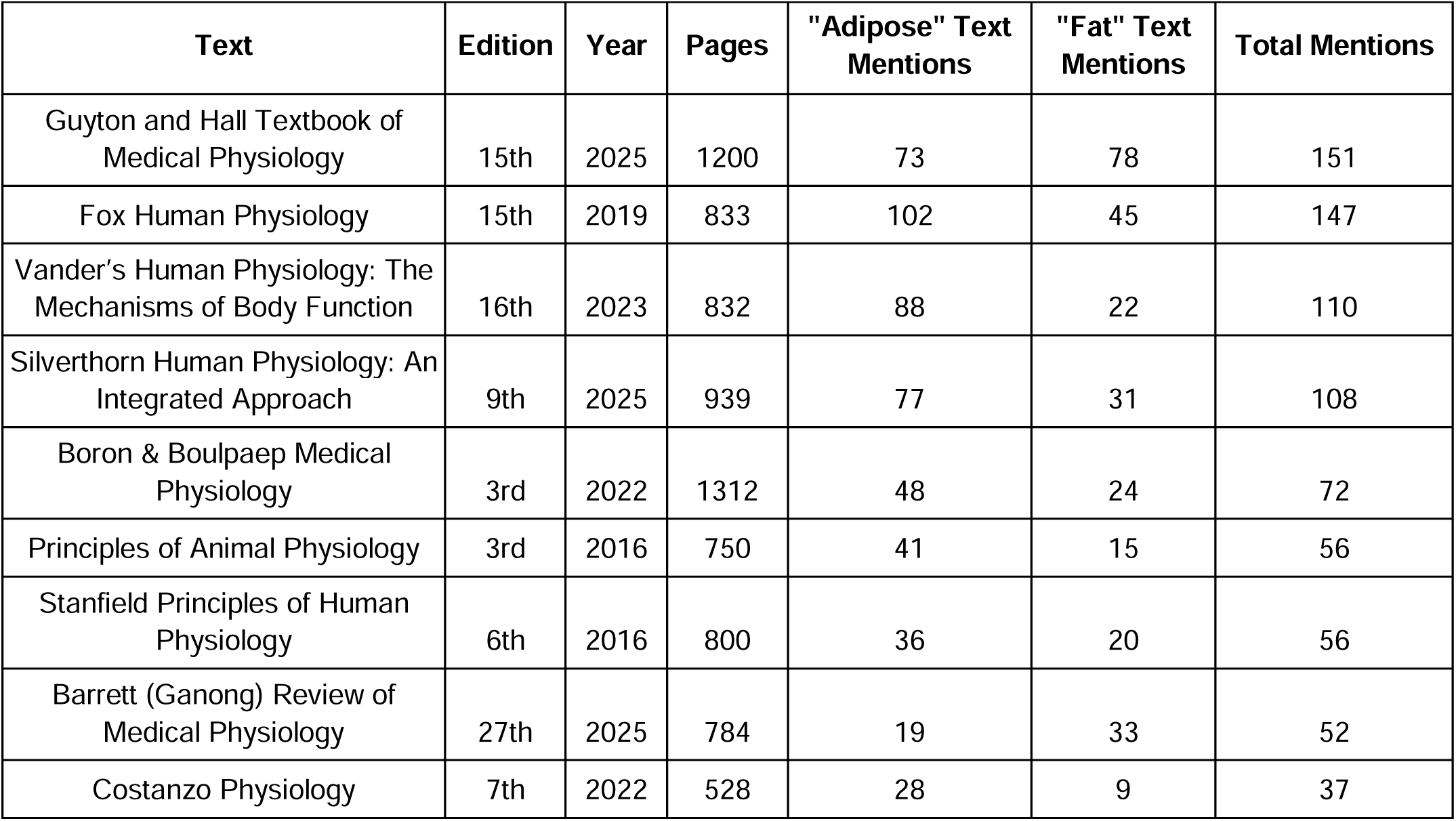
Physiology textbook terminology analysis. Texts are listed in descending order from highest number of adipose/fat mentions to the lowest number. “Total mentions” represents the combined count of both terms. Page numbers are provided for reference to overall textbook length.

| Text | Edition | Year | Pages | "Adipose" Text Mentions | "Fat" Text Mentions | Total Mentions |
| --- | --- | --- | --- | --- | --- | --- |
| Guyton and Hall Textbook of Medical Physiology | 15th | 2025 | 1200 | 73 | 78 | 151 |
| Fox Human Physiology | 15th | 2019 | 833 | 102 | 45 | 147 |
| Vander's Human Physiology: The Mechanisms of Body Function | 16th | 2023 | 832 | 88 | 22 | 110 |
| Silverthorn Human Physiology: An Integrated Approach | 9th | 2025 | 939 | 77 | 31 | 108 |
| Boron & Boulpaep Medical Physiology | 3rd | 2022 | 1312 | 48 | 24 | 72 |
| Principles of Animal Physiology | 3rd | 2016 | 750 | 41 | 15 | 56 |
| Stanfield Principles of Human Physiology | 6th | 2016 | 800 | 36 | 20 | 56 |
| Barrett (Ganong) Review of Medical Physiology | 27th | 2025 | 784 | 19 | 33 | 52 |
| Costanzo Physiology | 7th | 2022 | 528 | 28 | 9 | 37 |

Across textbooks, the term “fat” was significantly more likely than “adipose” to be used in indirect/non-functional (Figure 2B; OR = 1.95, p_adj = 0.0016) and obesity/excess contexts (OR = 1.90, p_adj = 0.0033). In contrast, “fat” was less likely to be used in metabolic contexts (OR = 0.48, p_adj < 0.001), indicating that mentions describing metabolic functions of adipose tissue more frequently use the term “adipose.” Differences in thermogenic, endocrine, and immune contexts were not statistically significant after correction for multiple comparisons. These patterns indicate that while both terms are used throughout physiology texts, “adipose” is more commonly associated with functional metabolic descriptions, whereas “fat” is more often used in indirect/non-functional or obesity-related contexts.

**Figure 2:**
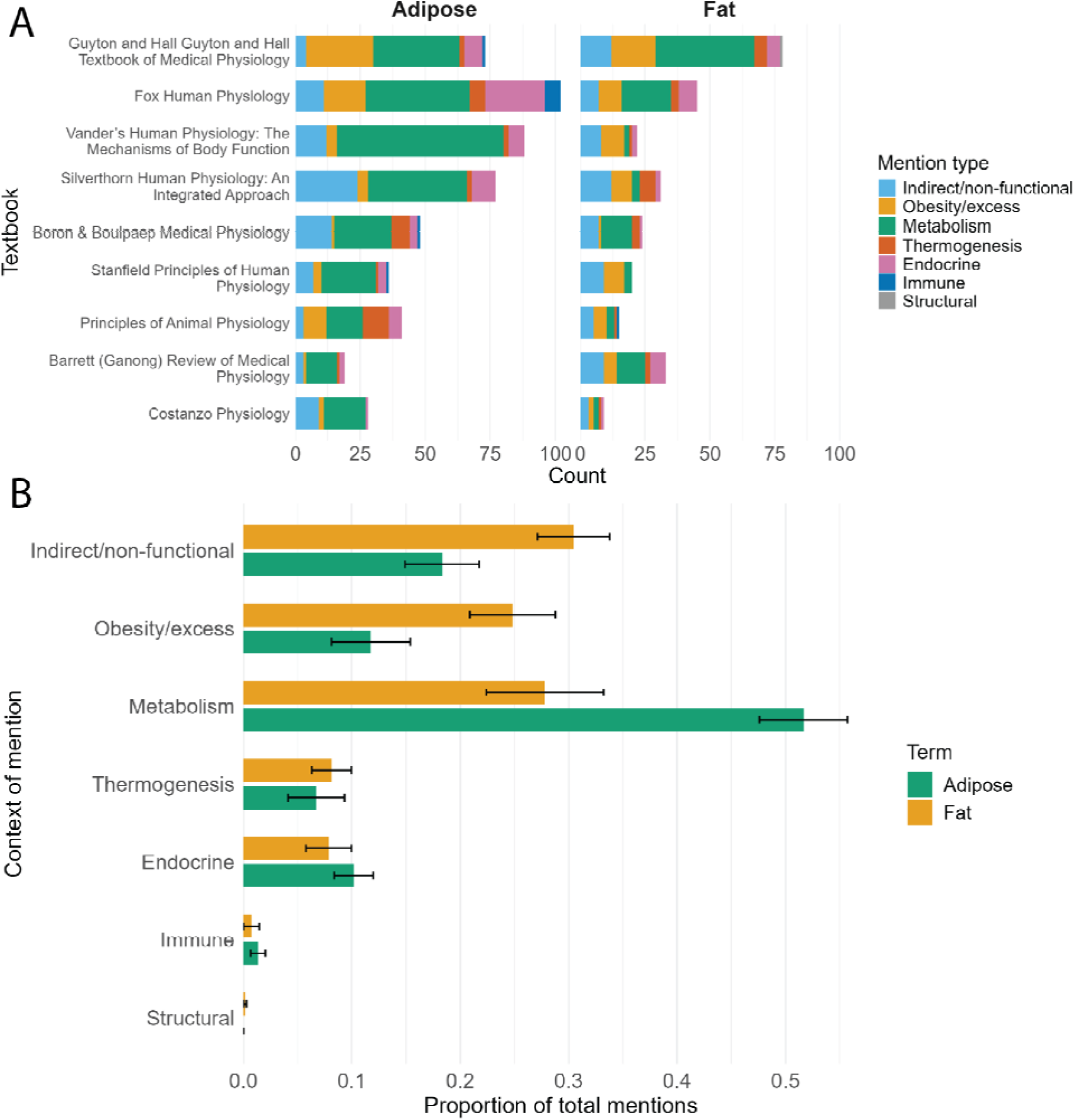
Terminology and contextual framing of adipose tissue in physiology textbooks. (A) Counts of “adipose” and “fat” mentions across textbooks, stratified by context of use (e.g., metabolic, endocrine, obesity-related). (B) Proportion of total mentions by context for each term (mean ± 95% CI).

Adipose tissue was infrequently depicted in physiology textbook images (Table S2, Figure S2). Across texts, adipose appeared in a small proportion of total images (typically <5%), and labeling rates were low, with depot-specific labeling nearly absent. Representation scores were skewed toward lower categories, with the largest proportion of images falling into score 1 (minimal representation), indicating that adipose tissue is often only partially depicted or shown incidentally in physiological figures.

### Anatomy & Physiology Textbooks

Adipose tissue was moderately represented in images across combined anatomy and physiology textbooks (Table 3, Figure 3). The proportion of images containing adipose ranged from 5.1% to 11.7% across texts. Despite this visual presence, labeling rates remained low, with typically fewer than 2% of total images including adipose labels and depot-specific labeling rare across all texts. Visual adipose representation scores were unevenly distributed (Figure 3B), with the largest proportion of images falling into score 1 (37.0%, 95% CI: 34.0–40.1), with lower but broadly similar proportions across higher categories (score 2: 20.8%, 95% CI: 18.4–23.4; score 3: 17.0%, 95% CI: 14.9–19.4; score 4: 25.2%, 95% CI: 22.5–28.2). This pattern indicates that adipose tissue is most often minimally represented, with fewer images occupying intermediate or more complete levels.

**Figure 3:**
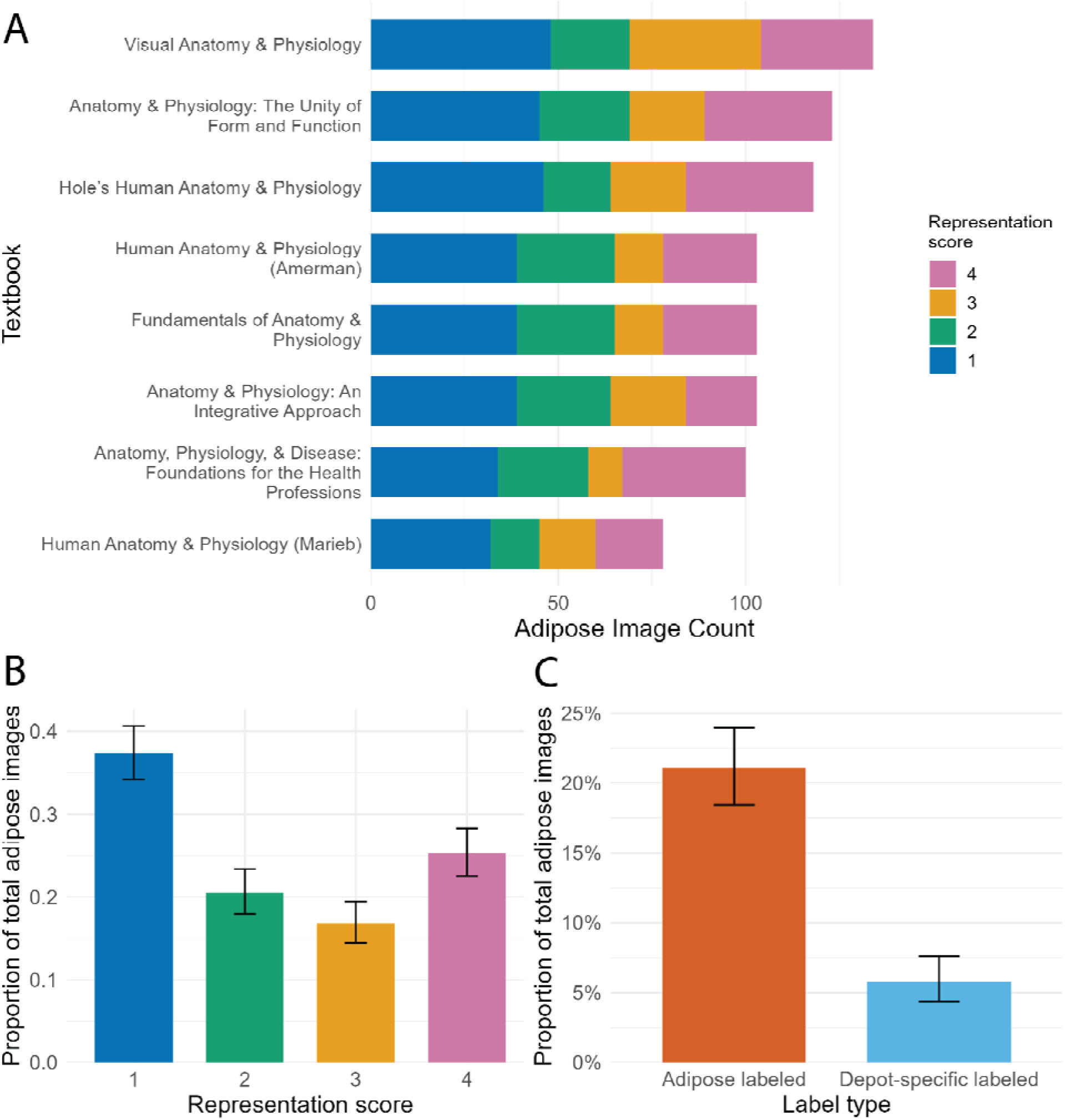
Adipose tissue representation and labeling in combined anatomy & physiology textbook images. (A) Counts of adipose-containing images by textbook, stratified by representation score (1–4), with higher scores indicating more complete depiction. (B) Proportion of adipose images in each representation category (mean ± 95% CI). (C) Proportion of images containing adipose with adipose labeling, including any label and depot-specific labels (mean ± 95% CI).

**Table 3:** Adipose tissue representation in combined anatomy & physiology textbook images. Texts are listed in descending order from highest proportion of visual adipose tissue representation to the lowest. Percentages are calculated relative to the total number of images within each textbook.

| Text | Edition | Year | Total images | Adipose images, n (%) | Images with adipose labels, n (%) | Depot-specific labels, n (%) |
| --- | --- | --- | --- | --- | --- | --- |
| Anatomy, Physiology, & Disease: Foundations for the Health Professions | 3rd | 2023 | 854 | 100 (11.7%) | 20 (2.3%) | 2 (0.2%) |
| Anatomy & Physiology: The Unity of Form and Function | 6th | 2012 | 1374 | 123 (9%) | 27 (2%) | 9 (0.7%) |
| Hole's Human Anatomy & Physiology | 15th | 2023 | 1380 | 118 (8.6%) | 17 (1.2%) | 5 (0.4%) |
| Fundamentals of Anatomy & Physiology | 12th | 2023 | 1432 | 103 (7.2%) | 26 (1.8%) | 8 (0.6%) |
| Human Anatomy & Physiology (Amerman) | 3rd | 2024 | 1510 | 103 (6.8%) | 26 (1.7%) | 8 (0.5%) |
| Visual Anatomy & Physiology | 3rd | 2018 | 2092 | 135 (6.5%) | 14 (0.7%) | 4 (0.2%) |
| Anatomy & Physiology: An Integrative Approach | 4th | 2022 | 1689 | 104 (6.2%) | 31 (1.8%) | 8 (0.5%) |
| Human Anatomy & Physiology (Marieb) | 11th | 2019 | 1515 | 78 (5.1%) | 21 (1.4%) | 6 (0.4%) |

In-text mentions of adipose and fat also varied across combined anatomy and physiology textbooks (Table 4, Figure 4). Total mention counts ranged from 29 to 142 per text, with both terms used extensively across books. In contrast to anatomy-only texts, references to “adipose” were often as frequent as or more frequent than references to “fat,” though this pattern varied by textbook. Across contexts, both terms were most commonly used in indirect or non-functional descriptions, followed by metabolic and obesity-related contexts, with relatively fewer mentions of thermogenic, endocrine, structural, or immune roles. Contextual usage differed systematically between terms. “Fat” was significantly more likely than “adipose” to be used in obesity- or excess-related contexts (OR = 2.56, 95% CI: 1.56–4.18, BH-adjusted p < 0.001), whereas it was significantly less likely to be used in metabolic contexts (OR = 0.49, 95% CI: 0.34–0.68, BH-adjusted p < 0.001). No significant differences were observed between terms for indirect/anatomical, thermogenic, endocrine, or structural contexts after correction for multiple comparisons (all BH-adjusted p ≥ 0.25).

**Figure 4:**
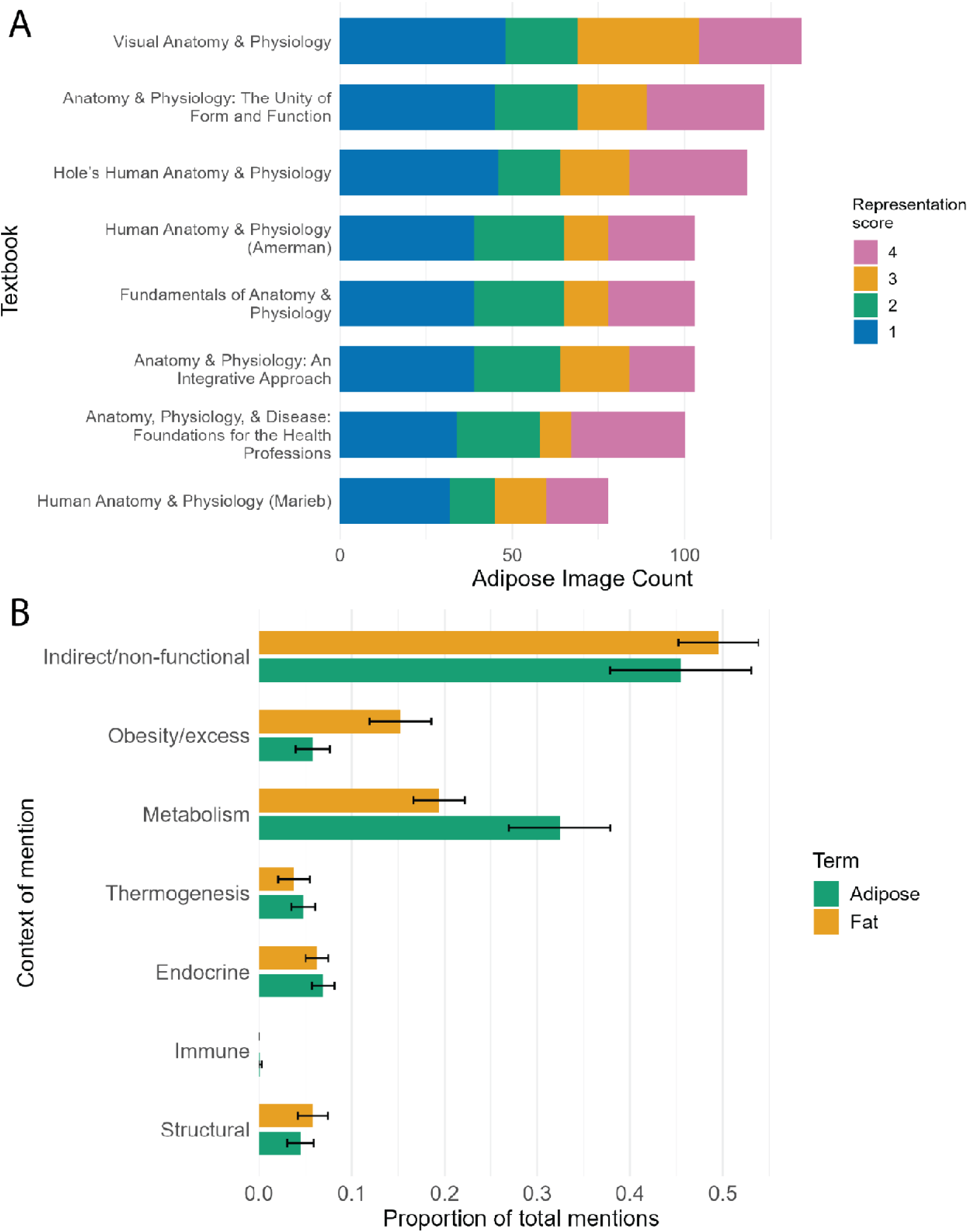
Terminology and contextual framing of adipose tissue in combined anatomy & physiology textbook images. (A) Counts of “adipose” and “fat” mentions across textbooks, stratified by context of use (e.g., metabolic, endocrine, obesity-related). (B) Proportion of total mentions by context for each term (mean ± 95% CI).

**Table 4:** Combined anatomy & physiology textbook terminology analysis. Texts are listed in descending order from highest number of adipose/fat mentions to the lowest number. “Total mentions” represents the combined count of both terms. Page numbers are provided for reference to overall textbook length.

| Text | Edition | Year | Pages | "Adipose" Text Mentions | "Fat" Text Mentions | Total Mentions |
| --- | --- | --- | --- | --- | --- | --- |
| Hole's Human Anatomy & Physiology | 15th | 2023 | 928 | 81 | 61 | 142 |
| Fundamentals of Anatomy & Physiology | 12th | 2023 | 1172 | 47 | 72 | 119 |
| Human Anatomy & Physiology (Marieb) | 11th | 2019 | 1137 | 76 | 29 | 105 |
| Human Anatomy & Physiology (Amerman) | 3rd | 2024 | 1116 | 54 | 50 | 104 |
| Anatomy, Physiology, & Disease: Foundations for the Health Professions | 3rd | 2023 | 663 | 39 | 60 | 99 |
| Anatomy & Physiology: The Unity of Form and Function | 6th | 2012 | 1136 | 70 | 26 | 96 |
| Visual Anatomy & Physiology | 3rd | 2018 | 1073 | 45 | 31 | 76 |
| Anatomy & Physiology: An Integrative Approach | 4th | 2022 | 1179 | 14 | 15 | 29 |

Overall, while combined anatomy and physiology texts are more likely than anatomy-only texts to use the term “adipose” and to describe its metabolic roles, terminology remains context-dependent: “fat” is preferentially used in excess-related framing, whereas “adipose” is more strongly associated with functional, physiological descriptions. Mentions nonetheless still occur most commonly in non-functional contexts.

## Discussion

Across anatomy, physiology, and combined anatomy and physiology textbooks, adipose tissue is consistently present but not consistently presented. Rather than being integrated as a coherent organ system, it is fragmented across visual and textual domains. It is commonly visible in images yet rarely labeled, described using inconsistent terminology, and unevenly associated with its known physiological functions. These findings point to a gap between representation and understanding, in which adipose tissue is present but not fully established as a distinct anatomical and functional system.

In anatomy textbooks, adipose tissue frequently appears in images, often in substantial quantities, yet is rarely labeled. This pattern is consistent across atlases, clinical texts, and general textbooks, indicating that under-labeling reflects a broader convention in anatomical representation rather than differences between resource types. The bimodal distribution of representation shows that images tend toward two extremes (minimal vs. extensive depiction). Images in which adipose tissue was fully represented were typically anatomical plane views (e.g., transverse or sagittal sections) or images from living individuals (e.g., MRI, dual-contrast CT scans, or surgical photographs), contexts in which adipose is unavoidable. Within artistic representations, a similar pattern was observed: images created using an anatomical donor as a direct reference or model tended to include more adipose tissue, whereas more schematic or concept-driven illustrations often minimized or omitted it. Even in these cases of higher visual representation, adipose tissue is often left unlabeled, limiting opportunities for students to recognize it as a defined anatomical component. Textual descriptions in anatomy texts reflect a similar pattern. Labeling and in-text mentions in anatomy textbooks predominantly use the term “fat” rather than “adipose.” This pattern may reflect broader standardization practices in anatomical nomenclature. Terminological resources such as *Terminologia Anatomica*, which guide naming conventions in anatomical education, include numerous Latin terms derived from adipos-, yet their English translations consistently default to “fat” (e.g., “fat pad,” “fat body”). This effectively positions “fat” as the guided term, although “adipose” more precisely reflects the complexity of the tissue. The term “fat,” by contrast, is imprecise: it may refer to lipids rather than to a tissue, and it reduces a heterogeneous, multicellular organ system to a single stored component. Anatomy texts may thus be reinforcing established nomenclature conventions rather than reflecting current understandings of adipose tissue.

In contrast, physiology textbooks use the term “adipose” at a higher or equivalent rate as “fat”, and commonly frame adipose tissue in functional terms, particularly in metabolic contexts. However, this functional framing is not comprehensive across adipose tissue’s known roles. Endocrine and thermogenic roles are consistently described across texts, whereas immune functions of adipose tissue are rarely mentioned, and structural roles are largely absent. This omission may, in part, reflect differences in pedagogical accessibility: endocrine functions, often centered on relatively straightforward signaling pathways such as leptin, lend themselves to simplified, canonical narratives. The known immune functions of adipose tissue are comparatively complex, with context-dependent interactions across multiple cell types and signaling networks that are less easily summarized for introductory materials. Structural roles may be similarly underrepresented as they do not fit as clearly within the regulatory, systems-focused emphasis of physiology texts and are often treated as background anatomical context rather than explicit functions. This functional emphasis is also reflected in patterns of terminology, as they are more likely to use the term “adipose” rather than “fat” in these contexts. Though, the term “fat” remains more strongly associated with indirect or obesity-related contexts, while “adipose” is preferentially used when describing metabolic processes. This divergence suggests that language itself may cue different conceptual framings, positioning ‘fat’ as passive or non-functional tissue and ‘adipose’ as an active, physiological entity. Despite this more functional emphasis in text, adipose tissue remains visually underrepresented in physiology textbooks, where it appears in relatively few images and is typically only partially depicted. This creates a mismatch between textual emphasis and visual representation, in which adipose tissue is discussed as physiologically important but is not consistently shown as a discrete anatomical entity.

Combined anatomy and physiology textbooks occupy an intermediate position. These texts are more likely than anatomy-only resources to use the term “adipose” and to describe its roles in metabolism, endocrine signaling, and thermogenesis, reflecting patterns observed in physiology texts, and are also more likely than other text types to include its structural functions. However, they retain limitations from both domains. Image-based representation remains constrained, with low labeling rates and a predominance of minimally depicted adipose tissue. Terminology patterns also persist, including continued preferential use of “fat” in non-functional or obesity-related contexts. Thus, even in combined anatomy and physiology curricula, adipose tissue is not consistently presented as a cohesive anatomical and physiological system.

Taken together, these findings suggest that adipose tissue is not absent from educational materials but is instead inconsistently and fragmentarily framed across visual and textual domains. Across text types, it may appear as some combination of the following: visible in images yet frequently unlabeled, described but not functionally emphasized, described using inconsistent terminology, unevenly associated with its functional roles, and functionally discussed without corresponding visual reinforcement. Importantly, these elements do not align to produce a cohesive representation. This fragmentation may therefore limit students’ ability to recognize adipose tissue as an integrated, multifunctional organ system. These patterns likely reflect broader historical and pedagogical conventions. Adipose tissue is typically classified as a connective tissue, a category that encompasses a wide range of structures with diverse functions. However, unlike other connective tissues such as bone, which are consistently labeled and emphasized, adipose tissue may be deprioritized due to its perceived role as excess, pathologic, or expendable tissue. As a result, adipose tissue may be recognized in classification but not consistently treated as a meaningful anatomical and physiological structure in its own right.

Additionally, in our analysis across all textbook types, the term “fat” was more frequently used in non-functional and obesity-related contexts, whereas “adipose” was preferentially used in functional physiological descriptions, indicating that terminology choice is not neutral. This patterned preferential use of “fat” may reinforce a framing of the tissue as passive or excessive, in contrast to “adipose,” which is more consistently aligned with active biological roles. This distinction carries broader linguistic and social implications. The term “fat” is not only a descriptor of biological tissue but also a socially charged term frequently used in stigmatizing ways (Martin-Wagar et al., 2024; Meadows & Daníelsdóttir, 2016; Rook & Holmes, 2023). Its use within educational materials, especially when paired with non-functional or pathological framing, may blur the boundary between anatomical description and socially embedded judgment, reinforcing negative or reductive associations. Within anatomy education, where adipose tissue is often under-labeled or treated as background rather than as a structure of interest, this terminology may further reinforce its marginalization. These patterns are particularly relevant given that anatomy education has been identified as a setting in which weight-related bias can be reproduced or reinforced (Goss et al., 2020). Within this environment, the combined effects of terminology and framing may shape not only how adipose tissue is conceptualized, but also how bodies characterized by greater adiposity are implicitly interpreted, with potential downstream implications for clinical perception and care (Puhl, 2023).

From an educational perspective, the inconsistent representation of adipose tissue across modalities may contribute to gaps in anatomical understanding. If adipose tissue is not labeled in images, students will not learn where it is expected to appear anatomically; if its functions are unevenly emphasized, with stronger coverage in metabolic contexts than in its broader endocrine, immune, thermogenic, and structural roles, its full physiological significance may be underappreciated. Improving the consistency of adipose representation, particularly through more explicit labeling and integration of structure and function, would support a more complete understanding of its biological importance.

Several limitations should be considered. This analysis was limited to a selected set of widely used textbooks and may not capture the full range of contemporary educational materials, particularly as many courses increasingly rely on instructor-curated resources such as primary literature, excerpts, and composite materials. Coding of representation and context, while systematic, involved some degree of subjectivity. In particular, the study does not establish a normative benchmark for how much adipose tissue *should* be represented in either images or text. Because anatomical figures are often designed to emphasize specific systems, and physiology content varies widely in scope, the expected inclusion and prominence of adipose tissue is inherently context-dependent. As a result, this study cannot assess underrepresentation in absolute terms, but instead characterizes patterns in how adipose tissue is depicted, labeled, and discussed when it is included. Finally, this study evaluates representation within educational materials rather than how students interpret or learn from them.

## Conclusion

Overall, these findings highlight a consistent pattern across educational resources: adipose tissue is present but not holistically integrated into anatomical and physiological frameworks. In anatomy texts, adipose tissue is present visually but not engaged; in physiology texts, it is discussed but not visualized; and in combined texts, it occupies an intermediate, incomplete position. This pattern is further reinforced by language, as the term ‘fat’ is often used in place of ‘adipose tissue,’ especially in non-functional or obesity-related contexts, potentially introducing stigmatizing connotations that influence how the tissue is perceived. Addressing these gaps is necessary to align educational representations of adipose tissue with its established roles as a dynamic organ system and to support a more accurate and integrated understanding within medical and biological education.

## References

Ahima, R.S. and Flier, J.S., 2000. Adipose tissue as an endocrine organ. Trends in Endocrinology & Metabolism, 11(8), pp.327–332.

Bagchi, D.P. and MacDougald, O.A., 2019. Identification and dissection of diverse mouse adipose depots. Journal of visualized experiments: JoVE, (149), pp.10–3791.

Beresheim, A., Zepeda, D., Pharel, M., Soy, T., Wilson, A.B. and Ferrigno, C., 2024. Anatomy’s missing faces: an assessment of representation gaps in atlas and textbook imagery. Anatomical Sciences Education, 17(5), pp.1055–1070.

Berry, D. C., Stenesen, D., Zeve, D., & Graff, J. M. (2013). The developmental origins of adipose tissue. Development, 140(19), 3939–3949.

Blüher, M. (2019). Obesity: global epidemiology and pathogenesis. Nature reviews endocrinology, 15(5), 288–298.

Corvera, S. (2021a). Adipose tissue: from amorphous filler to metabolic mastermind. The Biochemist, 43(2), 16–20.

Corvera, S. (2021b). Cellular heterogeneity in adipose tissues. Annual review of physiology, 83(1), 257–278.

Cypess, A. M. (2022). Reassessing human adipose tissue. New England Journal of Medicine, 386(8), 768–779.

Daley, S. F., Ali, M. A., Ohnuma, T., & Adigun, R. (2025). Anorexia and cachexia. In StatPearls [Internet]. StatPearls Publishing.

Devlin, M. J. (2021). Brown adipose tissue, nonshivering thermogenesis, and energy availability. In Evolutionary cell processes in primates (pp. 131-160). CRC Press.

Dragoo, J. L., Shapiro, S. A., Bradsell, H., & Frank, R. M. (2021). The essential roles of human adipose tissue: Metabolic, thermoregulatory, cellular, and paracrine effects. Journal of Cartilage & Joint Preservation, 1(3), 100023.

Emont, M. P., & Rosen, E. D. (2023). Exploring the heterogeneity of white adipose tissue in mouse and man. Current opinion in genetics & development, 80, 102045.

Fox, C. S., Massaro, J. M., Hoffmann, U., Pou, K. M., Maurovich-Horvat, P., Liu, C. Y., … & O’Donnell, C. J. (2007). Abdominal visceral and subcutaneous adipose tissue compartments: association with metabolic risk factors in the Framingham Heart Study. Circulation, 116(1), 39–48.

Frühbeck, G., Gómez-Ambrosi, J., Muruzábal, F. J., & Burrell, M. A. (2001). The adipocyte: a model for integration of endocrine and metabolic signaling in energy metabolism regulation. American Journal of Physiology-Endocrinology and Metabolism, 280(6), E827–E847.

Goss, A. L., Rethy, L., Pearl, R. L., & DeLisser, H. M. (2020). The “difficult” cadaver: weight bias in the gross anatomy lab. Medical education online, 25(1), 1742966.

Grant, R. W., & Dixit, V. D. (2015). Adipose tissue as an immunological organ. Obesity, 23(3), 512–518.

Harleen, A., Forer, R., Etsell, K., Mackey, C., Cherry-Bukowiec, J.R. and Sonneville, K.R., 2026. Addressing Weight Stigma in Medical Education: Insights and Strategies From a Nutrition Curriculum Review. Journal of Medical Education and Curricular Development, 13, p.23821205261420878.

Harvey, I., Boudreau, A., & Stephens, J. M. (2020). Adipose tissue in health and disease. Open biology, 10(12).

Heidebrecht, C., Fierheller, D., Martel, S., Andrews, A., Hollahan, A., Griffin, L., Meerai, S., Lock, R., Nabavian, H., D’Silva, C. and Friedman, M., 2024. Raising awareness of anti-fat stigma in healthcare through lived experience education: a continuing professional development pilot study. BMC Medical Education, 24(1), p.64.

Ibrahim, M. M. (2010). Subcutaneous and visceral adipose tissue: structural and functional differences. Obesity reviews, 11(1), 11–18.

Kahn, C. R., Wang, G., & Lee, K. Y. (2019). Altered adipose tissue and adipocyte function in the pathogenesis of metabolic syndrome. The Journal of clinical investigation, 129(10), 3990–4000.

Kamrani, P., Marston, G., & Jan, A. (2023). Anatomy, connective tissue. In StatPearls [Internet]. StatPearls Publishing.

Kershaw, E. E., & Flier, J. S. (2004). Adipose tissue as an endocrine organ. The Journal of Clinical Endocrinology & Metabolism, 89(6), 2548–2556.

Labusca, L., & Zugun-Eloae, F. (2018). The unexplored role of intra-articular adipose tissue in the homeostasis and pathology of articular joints. Frontiers in veterinary science, 5, 35.

Lafontan, M., 2012. Historical perspectives in fat cell biology: the fat cell as a model for the investigation of hormonal and metabolic pathways. American Journal of Physiology-Cell Physiology, 302(2), pp.C327–C359.

Lee, M. J., Wu, Y., & Fried, S. K. (2013). Adipose tissue heterogeneity: implication of depot differences in adipose tissue for obesity complications. Molecular aspects of medicine, 34(1), 1–11.

Li, Z., Bowers, E., Zhu, J., Yu, H., Hardij, J., Bagchi, D. P., … & MacDougald, O. A. (2022). Lipolysis of bone marrow adipocytes is required to fuel bone and the marrow niche during energy deficits. elife, 11, e78496.

Loft, A., Emont, M. P., Weinstock, A., Divoux, A., Ghosh, A., Wagner, A., … & Rosen, E. D. (2025). Towards a consensus atlas of human and mouse adipose tissue at single-cell resolution. Nature metabolism, 7(5), 875–894.

Luong, Q., Huang, J., & Lee, K. Y. (2019). Deciphering white adipose tissue heterogeneity. Biology, 8(2), 23.

Macdougall, C. E., Wood, E. G., Loschko, J., Scagliotti, V., Cassidy, F. C., Robinson, M. E., … & Longhi, M. P. (2018). Visceral adipose tissue immune homeostasis is regulated by the crosstalk between adipocytes and dendritic cell subsets. Cell metabolism, 27(3), 588–601.

Martin-Wagar, C.A., Melcher, K.A., Attaway, S.E., Bennett, B.L., Thompson, C.J., Kronenberger, O. and Penwell, T.E., 2024. Does terminology matter when measuring stigmatizing attitudes about weight? Validation of a brief, modified attitudes toward obese persons scale. Obesity Science & Practice, 10(4), p.e70005.

Maung, J.N., Schill, R.L., Nishii, A., Foss de Freitas, M., Obua, B.N., Nygård, M., Mendez-Casillas, M.D., Hermsmeyer, I.D., Gilio, D., Besci, O. and Chen, Y., 2025. Lamin A/C regulates lipid metabolism and inflammation: insights from models of familial partial lipodystrophy 2. The Journal of Clinical Investigation, pp.e198387–e198387.

Maung, J.N., Oral, E.A. and MacDougald, O.A., 2026. Lamin A/C in health, laminopathies, and familial partial lipodystrophy 2. Trends in Endocrinology & Metabolism.

Meadows, A. and Daníelsdóttir, S., 2016. What’s in a word? On weight stigma and terminology. Frontiers in psychology, 7, p.1527.

Miricescu, D., Balan, D.G., Tulin, A., Stiru, O., Vacaroiu, I.A., Mihai, D.A., Popa, C.C., Enyedi, M., Nedelea, A.S., Nica, A.E. and Stefani, C., 2021. Impact of adipose tissue in chronic kidney disease development. Experimental and therapeutic medicine, 21(5), p.539.

Nutter, S., Eggerichs, L. A., Nagpal, T. S., Ramos Salas, X., Chin Chea, C., Saiful, S., … & Yusop, S. (2024). Changing the global obesity narrative to recognize and reduce weight stigma: a position statement from the World Obesity Federation. Obesity Reviews, 25(1), e13642.

Puhl, R.M., 2023. Weight stigma and barriers to effective obesity care. Gastroenterology Clinics, 52(2), pp.417–428.

Richard, A. J., White, U., Elks, C. M., & Stephens, J. M. (2020). Adipose tissue: physiology to metabolic dysfunction. Endotext.

Rui, L. (2017). Brown and beige adipose tissues in health and disease. Comprehensive Physiology, 7(4), 1281–1306.

Rook, E.D. and Holmes, K.J., 2023. How language shapes anti-fat bias: comparing the effects of disease and fat-rights framing. Frontiers in Communication, 8, p.1284074.

Sahni, S., Shiralkar, M., Mohamed, S., Carroll, R., Jung, B., Gaba, R., & Yazici, C. (2017). Superior mesenteric artery syndrome: the dark side of weight loss. Cureus, 9(11).

Scheller, E. L., Doucette, C. R., Learman, B. S., Cawthorn, W. P., Khandaker, S., Schell, B., … & MacDougald, O. A. (2015). Region-specific variation in the properties of skeletal adipocytes reveals regulated and constitutive marrow adipose tissues. Nature communications, 6(1), 7808.

Trayhurn, P., & Beattie, J. H. (2001). Physiological role of adipose tissue: white adipose tissue as an endocrine and secretory organ. Proceedings of the Nutrition Society, 60(3), 329–339.

Tseng, Y. H. (2023). Adipose tissue in communication: within and without. Nature Reviews Endocrinology, 19(2), 70–71.

Wajchenberg, B. L. (2000). Subcutaneous and visceral adipose tissue: their relation to the metabolic syndrome. Endocrine reviews, 21(6), 697–738.

Wronska, A., & Kmiec, Z. J. A. P. (2012). Structural and biochemical characteristics of various white adipose tissue depots. Acta physiologica, 205(2), 194–208.

Zwick, R. K., Guerrero-Juarez, C. F., Horsley, V., & Plikus, M. V. (2018). Anatomical, physiological, and functional diversity of adipose tissue. Cell metabolism, 27(1), 68–83.

