## Supplemental tables and figures for "The Unlabeled Organ: Adipose tissue and normativity in anatomy and physiology texts"

Supplementary Tables and Figures

Table S1: Anatomy textbook textbook terminology analysis. Texts are listed in descending order from highest number of adipose/fat mentions to the lowest number. “Total mentions” represents the combined count of both terms. Page numbers are provided for reference to overall textbook length.

| **Text** | **Edition** | **Year** | **Type** | **Pages** | **"Adipose" Text Mentions** | **"Fat" Text Mentions** | **Total Mentions** |
| --- | --- | --- | --- | --- | --- | --- | --- |
| Gray’s Anatomy: The Anatomical Basis of Clinical Practice | 43rd | 2026 | Textbook | 1559 | 176 | 185 | 361 |
| Moore’s Clinically Oriented Anatomy | 10th | 2026 | Clinical Textbook | 1200 | 5 | 103 | 108 |
| Tortora’s Principles of Anatomy | 15th | 2020 | Textbook | 998 | 46 | 47 | 93 |
| Abrahams’ and McMinn’s Clinical Atlas of Human Anatomy | 8th | 2020 | Clinical Atlas | 376 | 13 | 63 | 76 |
| Marieb’s Human Anatomy | 9th | 2020 | Textbook | 816 | 19 | 54 | 73 |
| Gray's Anatomy for Students | 5th | 2023 | Textbook | 1114 | 2 | 47 | 49 |
| Gosling’s Human Anatomy: Color Atlas and Textbook | 6th | 2017 | Atlas | 421 | 1 | 41 | 42 |
| Grant’s Atlas of Anatomy | 16th | 2017 | Atlas | 816 | 1 | 27 | 28 |
| The Big Picture: Gross Anatomy | 2nd | 2019 | Textbook | 496 | 10 | 13 | 23 |
| Sobotta Atlas of Human Anatomy | 16th | 2019 | Atlas | ‎1376 | 11 | 12 | 23 |
| Gilroy Atlas of Anatomy | 5th | 2025 | Atlas | 814 | 1 | 10 | 11 |
| Rohen’s Photographic Anatomy | 9th | 2021 | Atlas | 519 | 0 | 1 | 1 |
| Netter Atlas of Human Anatomy | 8th | 2023 | Atlas | 730 | 0 | 0 | 0 |

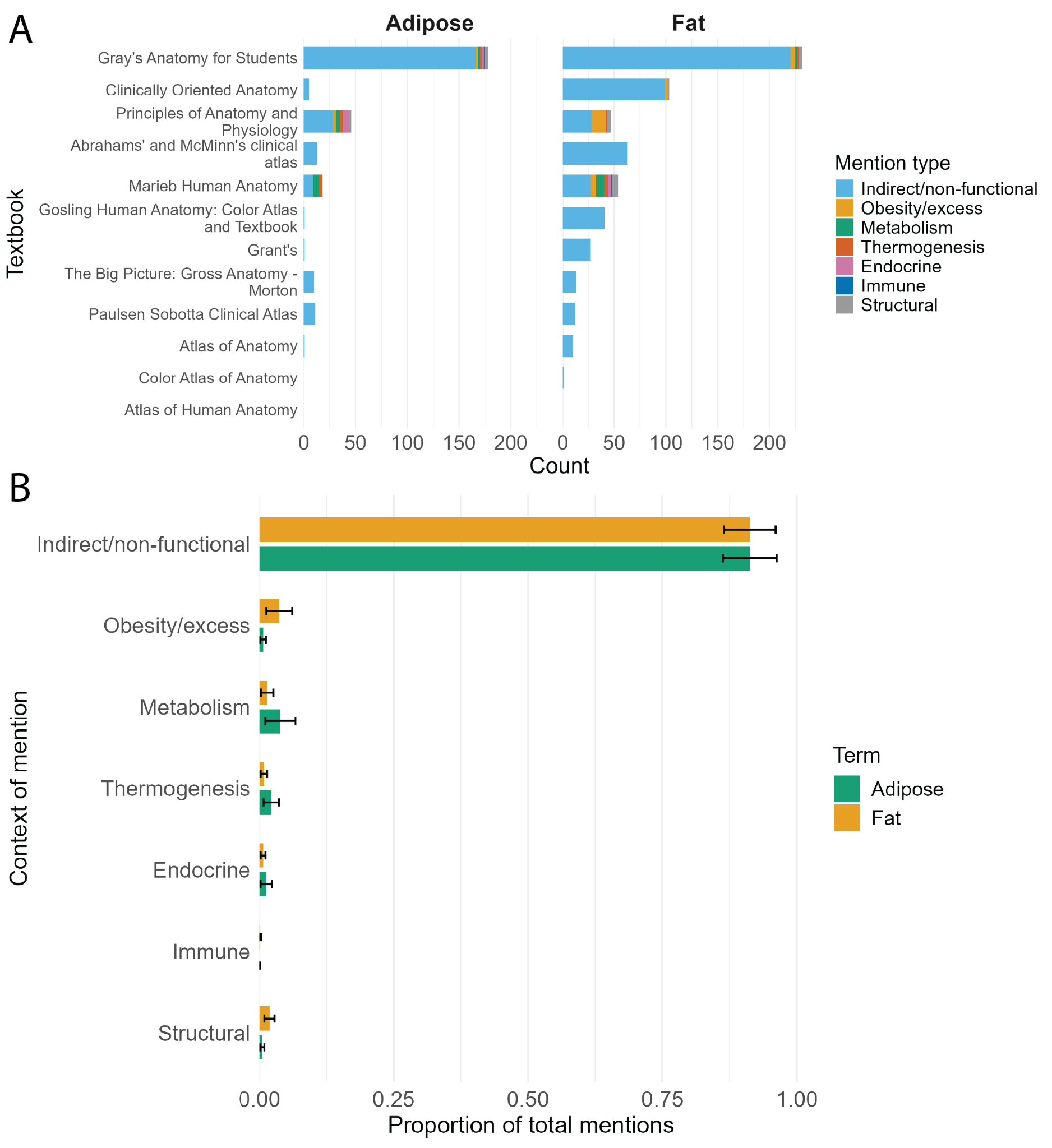

Fig S1: Terminology and contextual framing of adipose tissue in anatomy textbooks. (A) Counts of “adipose” and “fat” mentions across textbooks, stratified by context of use (e.g., metabolic, endocrine, obesity-related). (B) Proportion of total mentions by context for each term (mean ± 95% CI).

Table S2: Adipose tissue representation in physiology textbook images. Texts are listed in descending order from highest proportion of visual adipose tissue representation to the lowest. Percentages are calculated relative to the total number of images within each textbook.

| **Text** | **Edition** | **Year** | **Total images** | **Adipose images, n (%)** | **Images with adipose labels, n (%)** | **Depot-specific labels, n (%)** |
| --- | --- | --- | --- | --- | --- | --- |
| Guyton and Hall Textbook of Medical Physiology | 15th | 2025 | 909 | 12 (1.3%) | 6 (0.7%) | 1 (0.1%) |
| Fox Human Physiology | 15th | 2019 | 787 | 41 (5.2%) | 14 (1.8%) | 0 |
| Vander’s Human Physiology: The Mechanisms of Body Function | 16th | 2023 | 833 | 17 (2.0%) | 2 (0.24%) | 0 |
| Silverthorn Human Physiology: An Integrated Approach | 9th | 2025 | 1065 | 43 (4.0%) | 10 (0.94%) | 0 |
| Boron & Boulpaep Medical Physiology | 3rd | 2022 | 889 | 5 (0.6%) | 3 (0.3%) | 0 |
| Principles of Animal Physiology | 3rd | 2016 | 849 | 8 (0.94%) | 6 (0.71%) | 0 |
| Stanfield Principles of Human Physiology | 6th | 2016 | 1016 | 12 (1.2%) | 1 (0.1%) | 0 |
| Barrett (Ganong) Review of Medical Physiology | 27th | 2025 | 641 | 3 (0.47%) | 0 | 0 |
| Costanzo Physiology | 7th | 2022 | 343 | 2 (0.58%) | 2 (0.58%) | 0 |

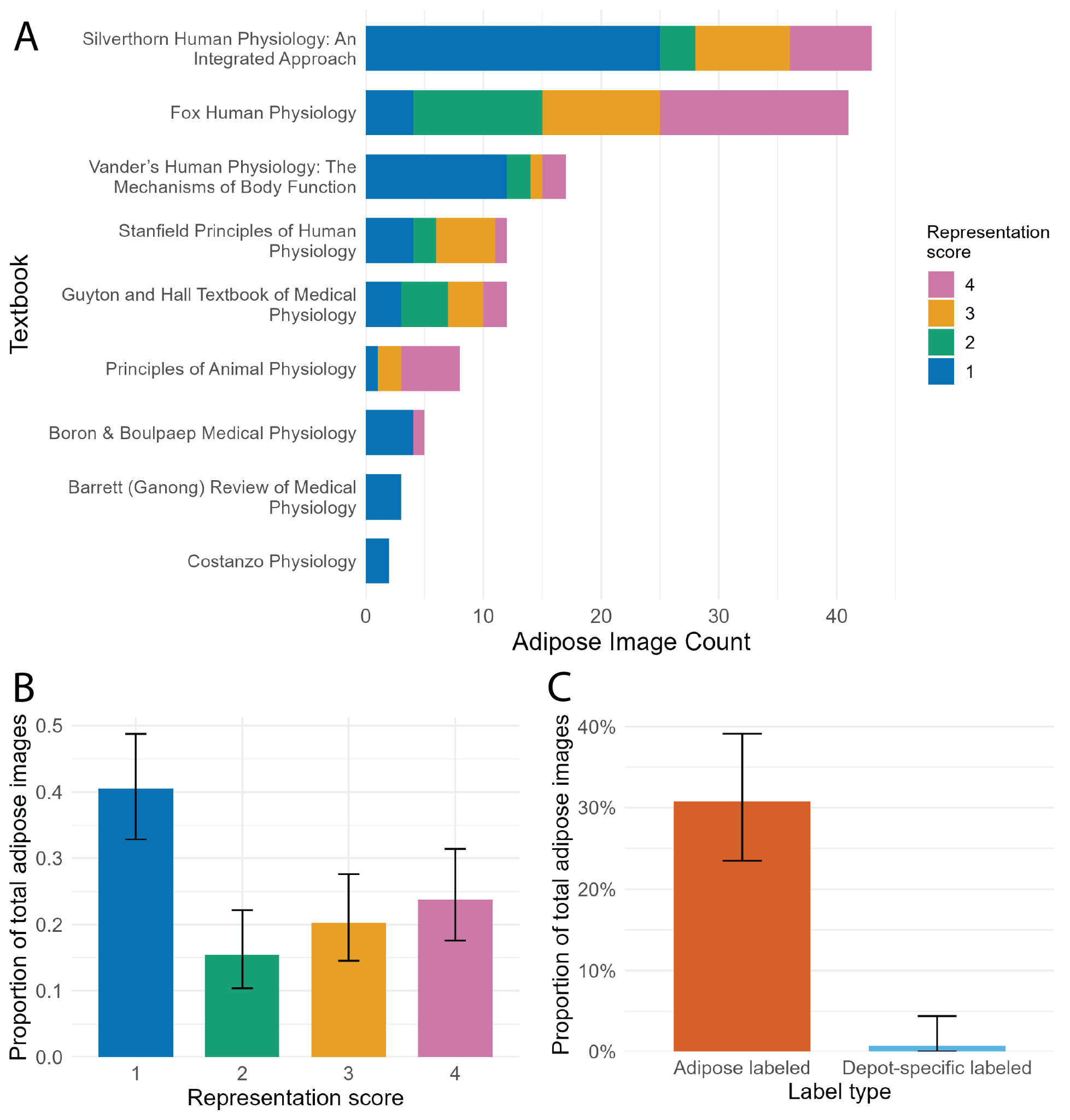

Fig S2: Terminology and contextual framing of adipose tissue in combined physiology textbook images. (A) Counts of “adipose” and “fat” mentions across textbooks, stratified by context of use (e.g., metabolic, endocrine, obesity-related). (B) Proportion of total mentions by context for each term (mean ± 95% CI).
